# Structural biologists, let’s mind our colors

**DOI:** 10.1101/2020.09.22.308593

**Authors:** Shyam M. Saladi, Ailiena O. Maggiolo, Kate Radford, William M. Clemons

## Abstract

In structural biology, most figures of macromolecules are aimed at those well-versed in structure, requiring prior familiarity with scales and commonly used color schemes. Yet, as structural biology becomes democratized with the increasing pace of structure determination, the accessibility of structural data is paramount. Here, we identify three keys, and have written accompanying software plugins, for structural biologists to create figures truer to the hard-won data and clearer across different modes of color vision and to non-expert readers.

1. Use perceptually uniform colormaps
2. Consider readers with different modes of color vision
3. Be explicit about scales and color usage

## Main Text

To visualize numerical relationships, we rely on colormaps to convert quantitative data into colors. Colormapping converts numerical data to fractional numbers between zero and one and then transforms the values to given colors, which can be visually represented using primary colors. Ideally, a colormap seeks to reproduce variation in data as perceived variation in color, yet widely used rainbow colormaps fail to do so. Instead, they introduce visual artefacts (Borland and Taylor, 2007; Moreland, 2016) that affect interpretation and can lead to critical diagnostic errors (Borkin et al., 2011). The most frequently used rainbow colormap, “Jet”, is formulated as a piecewise linear function of color primaries (Fig. 1a). These rainbow colormaps are problematic in two important ways. First, our brain perceives the variation in different parts of the color spectrum differently, which can be visualized for Jet by a jagged, irregular trace in the perceived difference (ΔE) for full-color vision (Fig. 1b, black line). Second, the irregularity in perceived difference becomes more pronounced for the 8% of the population with different modes of color vision (red-green and blue-green color blindness) (Fig. 1b).

**Figure 1:**
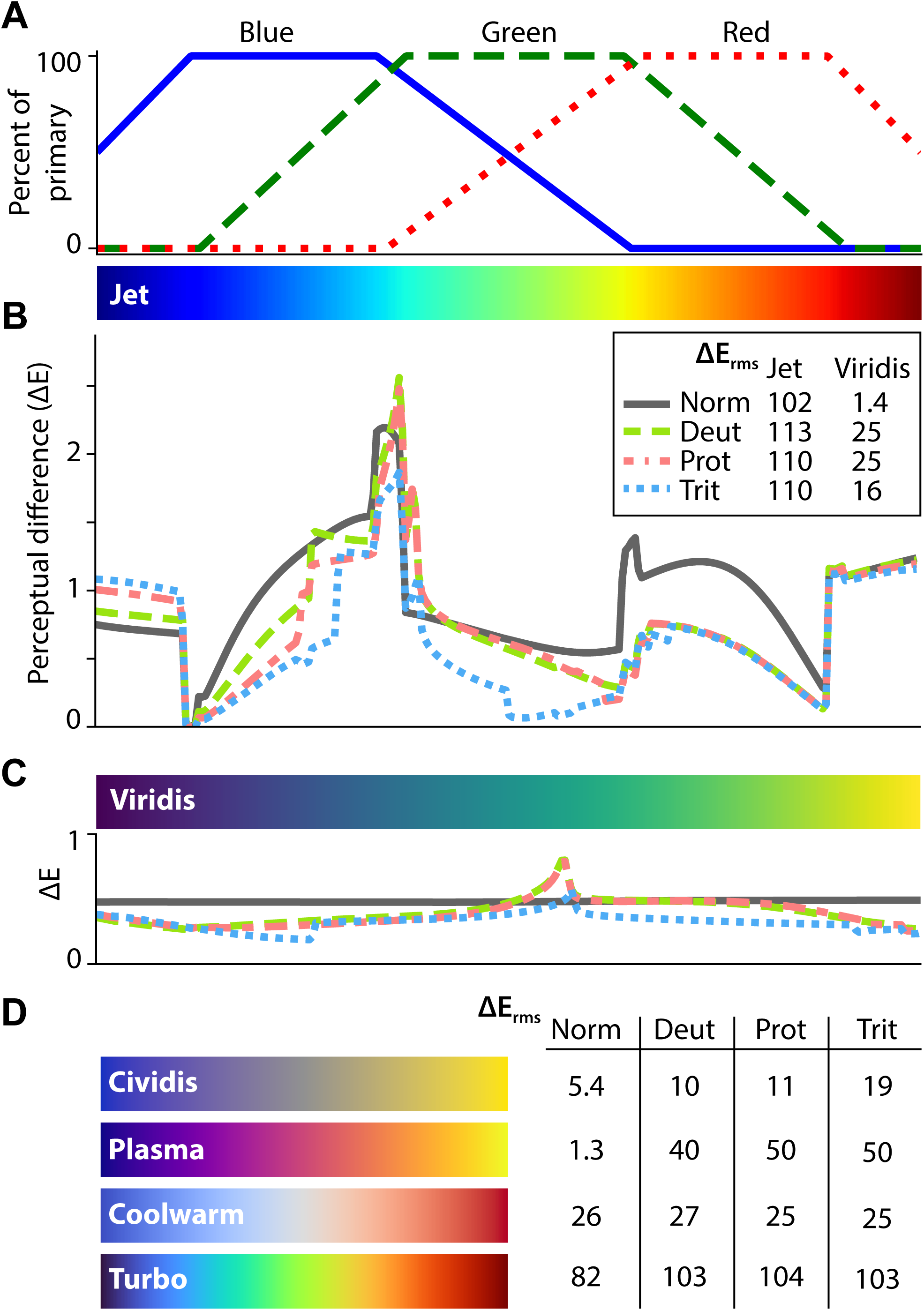
(a) The rainbow colormap Jet is constructed as a piecewise linear function of color primaries. Perceptual difference, ΔE, is the distance between two colors that reflects the ability to perceive a difference (CAM02-UCS, Moroney et al., 2002), of successive colors across (b) Jet and (c) Viridis under models of color vision: Norm, normal; Deut, deuteranopia (red-green color blind); Prot, protanopia (red-dichromacy); Trit, tritanopia (blue-yellow color blind). Note identically scaled y-axes. (b, inset) The root mean squared deviation in ΔE helps qualify how far each colormap is from ideal, *i.e*. constant ΔE. (d) A sample of performant colormaps: Cividis, engineered to be perceptually uniform across models of color vision deficiency; Plasma, perceptually uniform and among the series published with Viridis; Coolwarm, an efficient colormap for diverging data, *e.g*. electrostatic data where a natural midpoint exists; Turbo, an alternative to Jet where a rainbow colormap is obligatory, with better performance under normal vision but equally poor performance considering color vision deficiency.

Perceptually uniform colormaps present a clear solution by ensuring that the visually perceived differences match the underlying numerical difference in the data (Kovesi, 2015). Many such colormaps have already been engineered to trace smooth paths through perceptually uniform color space and, therefore, smoothly vary in perceived difference under various models of color vision (Smith and van der Walt, 2015; Nuñez et al., 2018). Looking closely at one such colormap, Viridis shows a more consistent ΔE across its colors, summarized by ΔE_rms_ values closer to zero compared to Jet (Fig. 1bc). However, improved colormaps, *e.g*. Viridis, Cividis, Plasma, are seldom used in structural analyses (Fig. 1d). To motivate broad adoption among structural biologists, we suggest adoption of “Viridis” and have implemented it in patches for PyMOL, ChimeraX, and VMD to facilitate uptake. The code is provided for the developers of these and other applications to broadly adopt.

An example of the improvement can be seen by looking at atomic B-factors in association with a structure of the SecY translocon (PDB: 1RH5, Van den Berg et al., 2004). Using a rainbow colormap, as typical (Fig. 2a), gives the false impression of a sharp transition in flexibility at the ends of the transmembrane alpha helices. This is not the case as apparent when viewed using Viridis (Fig. 2b). The distinction between the two types of colormaps becomes even more significant for readers with red-green color blindness. Jet results in a complete loss of all the information (Fig. 2c), while switching to Viridis accurately maintains interpretability for color blind readers (Fig. 2d).

**Figure 2:**
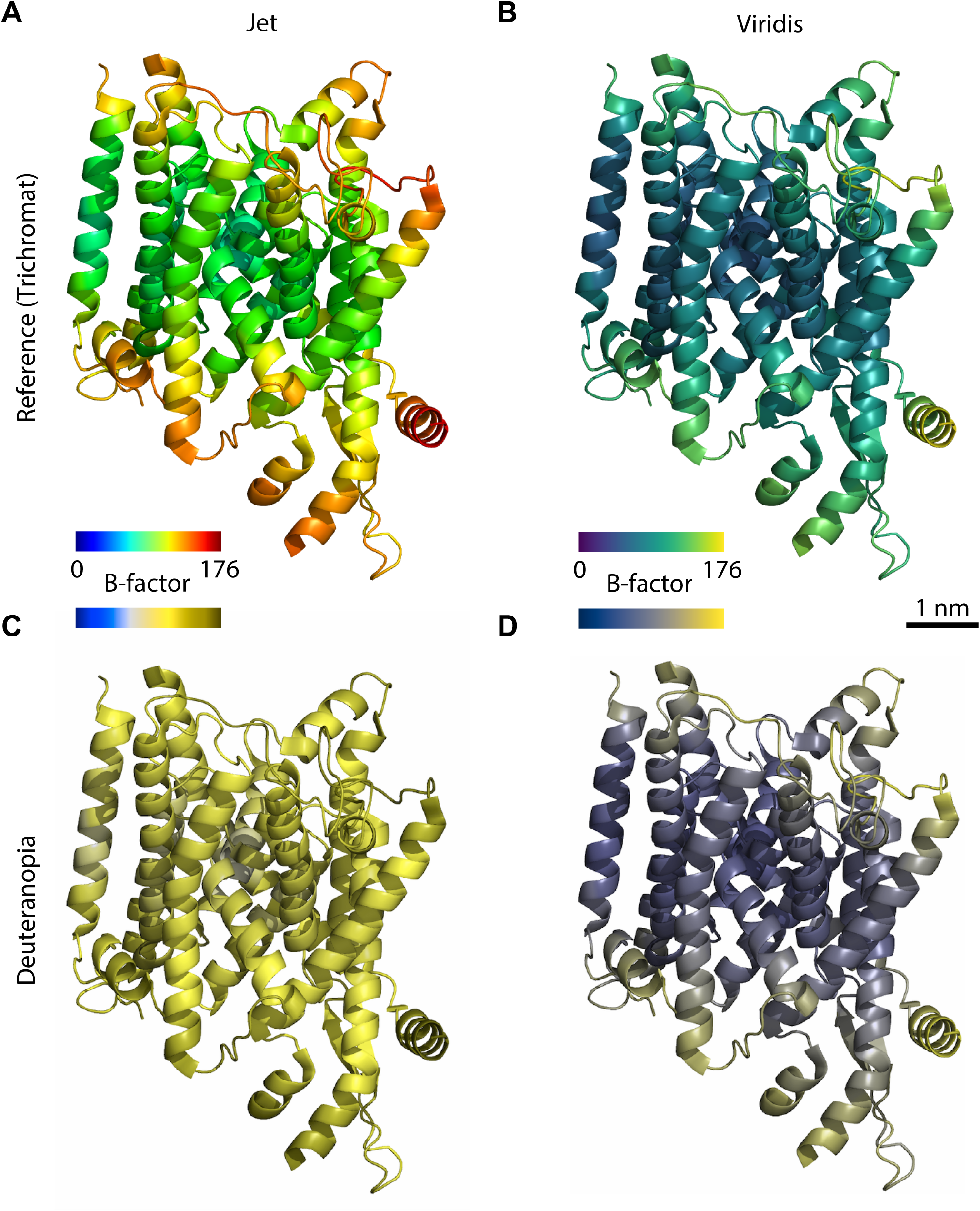
The SecY translocon (PDB: 1RH5, Van den Berg et al., 2004) colored by B-factor with the rainbow colormap Jet (a) suggests a sharp transition to flexibility at the extents of transmembrane alpha helices. The perceptually uniform colormap Viridis (b) provides a truer depiction showing that this transition is smooth. Under a model of complete deuteranopia, overlaid B-factor data is obscured when using Jet (c) but equally clear using Viridis (d). A colorbar indicating the mapping of B-factor to color and a scalebar is included.

With new colormaps and an increased focus on accessibility, we also owe it to readers to be explicit about the scale being represented. Scalebars will help interpretability for non-experts and easily double as a legend for the colormap used throughout a structural model (Fig. 2). This is especially critical considering our highly stylized representations of biological macromolecules (Richardson, 2000), the fact that ever-larger macromolecular complexes are being solved, and the quantitative nature of the data being presented, such as for tomographic densities. Colormaps, like Viridis, with scalebars are critical throughout experimental and computational structural biology, such as in displays of the directionality/orientation of polymers, local resolution, particle orientation bias, residue conservation, electrostatic potential, occupancy, radius of gyration, and others. We provide instructions to create scalebars here.

Similar to colormaps, figures with discrete colors require revisiting as these can be particularly problematic for different modes of color vision. A clear example are electron density difference (F_o_-F_c_) maps, visualized here for the P-cluster active site of nitrogenase with one cluster conformation removed (PDB: 3U7Q, Spatzal et al., 2011) (Fig. 3a). The maps clearly indicate two conformations of the cluster with differing occupancy. Using conventional green/red for the positive and negative difference maps, those with red-green deficiency cannot differentiate the two maps (Fig 3b). Moreover, since legends are rarely included, readers less familiar with structure analysis cannot easily recognize which color indicates density ‘modelled but missing’ versus ‘present but not modelled’. Moving to a different scheme, such as green/purple or blue/yellow, and including a key solves both issues (Fig. 2cd). Beautiful and color blind-friendly palettes are easy to create for all cases where discrete colors are needed (*e.g*. colorbrewer2.org, medialab.github.io/iwanthue, kuler.adobe.com, colorcyclesurvey.mpetroff.net, Okabe and Ito, 2002).

**Figure 3:**
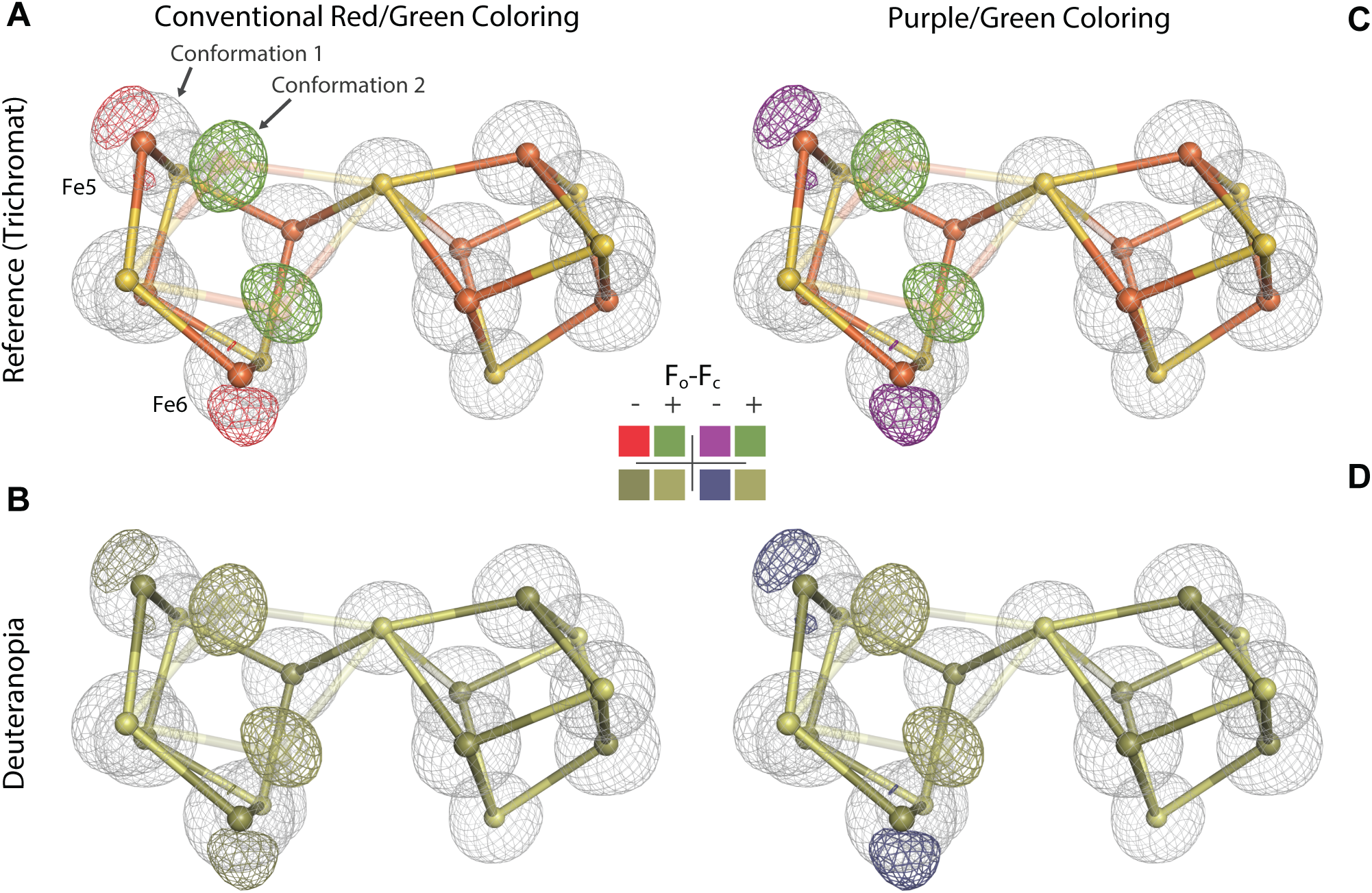
Electron density maps around the P-cluster of the nitrogenase MoFe protein with typical red/green coloring (PDB: 3U7Q, Spatzal et al., 2011). The 2F_o_-F_c_ map is shown as gray mesh and atoms of S and Fe are shown as yellow and orange spheres respectively. (a) The difference map (F_o_-F_c_) highlights the two conformations, *i.e*. positive/unmodelled density (+, green) where the second conformation is missing and negative/overmodelled density (−, red) where two Fe atoms are modelled at too high of an occupancy. Under a model of complete deuteranopia (*i.e*. red-green color blindness, b), this data is inaccessible. When using green and purple with a legend (c). the maps are clearly visualized under the model of deuteranopia (d). A key defining each color is included.

It is easy to either check colormaps for perceptual uniformity (Rogowitz and Kalvin, 2001) or figures for color blindness-accessibility (e.g. colororacle.org, [Colorblindly](https://github.com/oftheheadland/Colorblindly)). For programs where a non-ideal palette is default, users can implement patches, ask for direction on changing settings, or request developers rethink previous design decisions (Franklin, 2013; Sjores, 2020). This can be done by switching to a more optimal preset that would follow the lead of widely used scientific software suites, e.g. MATLAB, matplotlib (Droettboom, 2016; Eddins, 2014). For developers who prefer not to use preset color palettes, color transformation utilities are implemented in a variety of languages (Table S1).

We structural biologists are molecular cartographers. Just as mapmakers scrutinize each of their lines, symbols, and colors, it is incumbent upon us to represent hard-won snapshots of molecules both accessibly and faithfully to the underlying data.

## Methods

### Perceptual difference calculation

While there are numerous approaches to assess colormap uniformity and accessibility (Kovesi, 2015), here we have plotted the perceptual difference between each successive color within a colormap, which is the method by Smith and van der Walt, 2015. Colormaps are sourced from matplotlib as a series of 256 sRGB colors. In short, each color was converted from sRGB to the uniform colorspace CIECAM02-UCS through sRGB1-linear and XYZ100. ΔE is then the Euclidian difference between pairs of color coordinates. Perceptual differences under alternative modes of color vision were calculated by first finding the set of sRGB values that reproduce how each color of a colormap is perceived under alternative conditions (Machado et al., 2009).

All color space transformations were done through colorspacious (Smith et al., 2018). For further detail, see the Jupyter notebook used to create figures included with this manuscript.

### Structure visualizations

Atomic coordinates and/or electron density maps were downloaded from the RCSB Protein Data Bank (SecY accession 1RH5; nitrogenase MoFe protein accession 3U7Q). SecY was colored by B-factor using the default settings in PyMOL (`spectrum b`).

The coordinates of the nitrogenase MoFe protein contain two alternate conformations for the P-cluster. The atomic coordinates of one of the conformations of the cluster was deleted (molecule CLF in chain A) and the remaining conformation (molecule 1CL) was set to full occupancy and refined using REFMAC5 (Kovalevskiy et al., 2018). The refined density maps were converted to ccp4 maps using FFT within CCP4 (Ten Eyck, 1973; Winn et al., 2011).

Snapshots were generated using PyMOL v2.3.4 (Schrödinger, Inc., New York, NY) and rendered under deuteranopia using Color Oracle (Jenny, 2020) or colorspacious (Smith et al., 2018).

### Software availability

Patches to make Viridis and accompanying colormaps easily accessible in PyMOL, ChimeraX, and VMD can be found here: https://github.com/smsaladi/pymol_viridis, https://github.com/smsaladi/chimerax_viridis, https://github.com/smsaladi/vmd_viridis. A guide for creating a scalebar across various visualization programs can be found here: https://github.com/smsaladi/structure_scales. All code is made available under the open-source MIT License and will be deposited in the CaltechDATA institutional repository.

## Supporting information

Jupyter Notebook for Fig 1

## Acknowledgements

We thank members of the Clemons and Rees labs for discussion and S. Petrovic for comments on the manuscript. This work was supported by the National Institutes of Health (NIH) grants GM105385 and GM097572 (to WMC) and a National Science Foundation Graduate Research fellowship Grant 1144469 (to SMS).

---

**Table S1:**
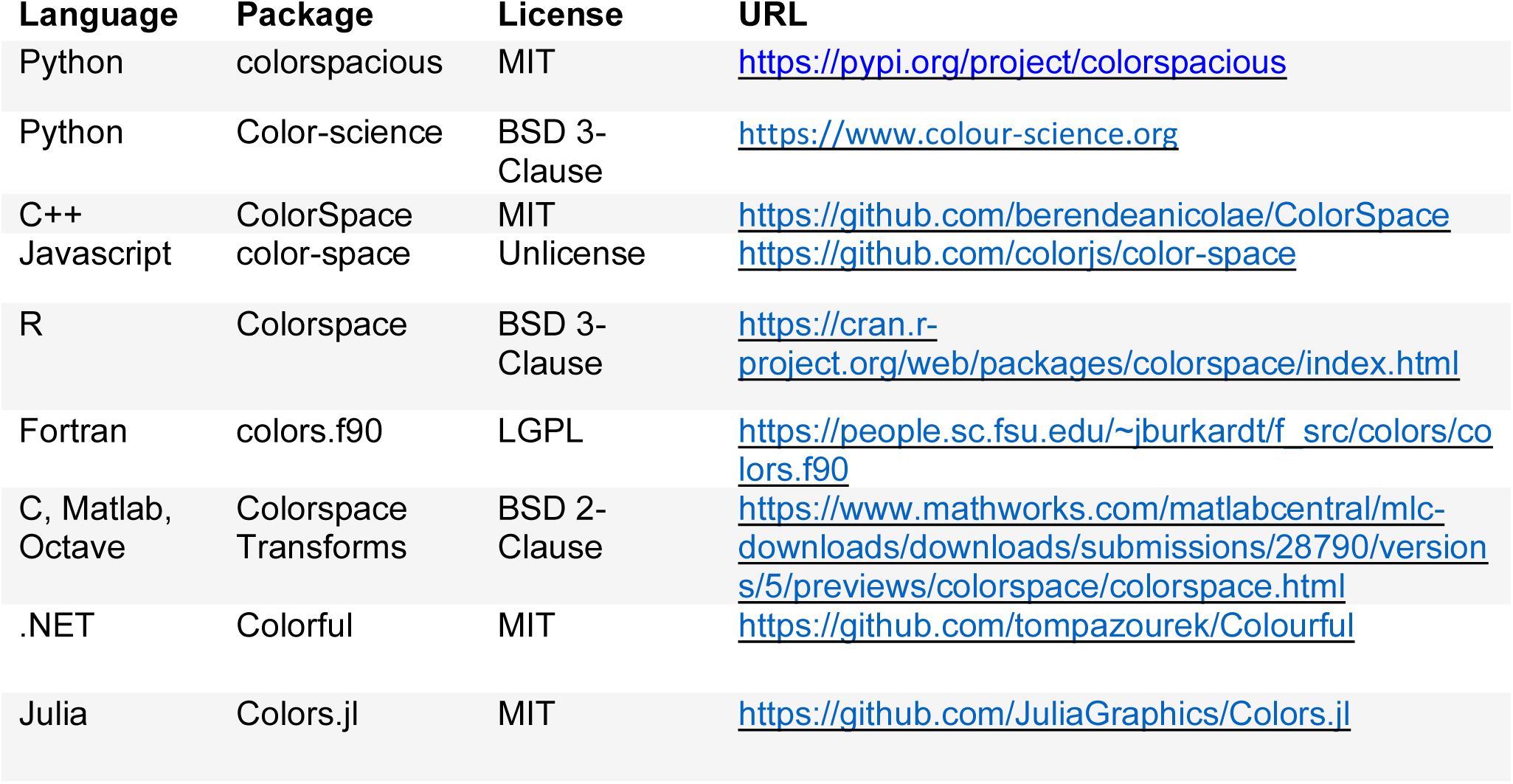
A non-exhaustive list of color space transformation codes implemented in various languages

