## Supplementary material for "Structural biologists, let’s mind our colors": Jupyter Notebook for Fig 1: colormap-discernibility.html


### Perceptual differences across colormaps¶

To accompany,

License: BSDv3

Date: September 2020

Parts adopted from Matthew Petroff are noted

In [1]:

```
import numpy as np

import matplotlib
import matplotlib.pyplot as plt

import colorspacious
import colorcet

matplotlib.rcParams['pdf.fonttype'] = 42
matplotlib.rcParams['ps.fonttype'] = 42
```

##### Register colormaps under CVD with matplotlib¶

In [2]:

```
def register_cvd(name, cvd_type):
    cmap = matplotlib.cm.get_cmap(name)(np.arange(256))[:, :3]
    cmap_cvd = colorspacious.cspace_convert(cmap, {
            "name": "sRGB1+CVD",
            "cvd_type": cvd_type,
            "severity": 100
        }, 'sRGB1')
    cmap_cvd = np.clip(cmap_cvd, 0, 1)
    name_cvd = name + '_' + cvd_type[:4]
    matplotlib.cm.register_cmap(name=name_cvd, cmap=matplotlib.colors.ListedColormap(cmap_cvd))
    
    print(name_cvd)
    return

register_cvd("viridis", "deuteranomaly")
register_cvd("jet", "deuteranomaly")
```

```
viridis_deut
jet_deut
```

##### Functions to calculate and plot Perceptual Differences across a colormap¶

In [3]:

```
# Turbo is from: https://gist.github.com/mikhailov-work/ee72ba4191942acecc03fe6da94fc73f
turbo_colormap_data = np.array([[0.18995,0.07176,0.23217],[0.19483,0.08339,0.26149],[0.19956,0.09498,0.29024],[0.20415,0.10652,0.31844],[0.20860,0.11802,0.34607],[0.21291,0.12947,0.37314],[0.21708,0.14087,0.39964],[0.22111,0.15223,0.42558],[0.22500,0.16354,0.45096],[0.22875,0.17481,0.47578],[0.23236,0.18603,0.50004],[0.23582,0.19720,0.52373],[0.23915,0.20833,0.54686],[0.24234,0.21941,0.56942],[0.24539,0.23044,0.59142],[0.24830,0.24143,0.61286],[0.25107,0.25237,0.63374],[0.25369,0.26327,0.65406],[0.25618,0.27412,0.67381],[0.25853,0.28492,0.69300],[0.26074,0.29568,0.71162],[0.26280,0.30639,0.72968],[0.26473,0.31706,0.74718],[0.26652,0.32768,0.76412],[0.26816,0.33825,0.78050],[0.26967,0.34878,0.79631],[0.27103,0.35926,0.81156],[0.27226,0.36970,0.82624],[0.27334,0.38008,0.84037],[0.27429,0.39043,0.85393],[0.27509,0.40072,0.86692],[0.27576,0.41097,0.87936],[0.27628,0.42118,0.89123],[0.27667,0.43134,0.90254],[0.27691,0.44145,0.91328],[0.27701,0.45152,0.92347],[0.27698,0.46153,0.93309],[0.27680,0.47151,0.94214],[0.27648,0.48144,0.95064],[0.27603,0.49132,0.95857],[0.27543,0.50115,0.96594],[0.27469,0.51094,0.97275],[0.27381,0.52069,0.97899],[0.27273,0.53040,0.98461],[0.27106,0.54015,0.98930],[0.26878,0.54995,0.99303],[0.26592,0.55979,0.99583],[0.26252,0.56967,0.99773],[0.25862,0.57958,0.99876],[0.25425,0.58950,0.99896],[0.24946,0.59943,0.99835],[0.24427,0.60937,0.99697],[0.23874,0.61931,0.99485],[0.23288,0.62923,0.99202],[0.22676,0.63913,0.98851],[0.22039,0.64901,0.98436],[0.21382,0.65886,0.97959],[0.20708,0.66866,0.97423],[0.20021,0.67842,0.96833],[0.19326,0.68812,0.96190],[0.18625,0.69775,0.95498],[0.17923,0.70732,0.94761],[0.17223,0.71680,0.93981],[0.16529,0.72620,0.93161],[0.15844,0.73551,0.92305],[0.15173,0.74472,0.91416],[0.14519,0.75381,0.90496],[0.13886,0.76279,0.89550],[0.13278,0.77165,0.88580],[0.12698,0.78037,0.87590],[0.12151,0.78896,0.86581],[0.11639,0.79740,0.85559],[0.11167,0.80569,0.84525],[0.10738,0.81381,0.83484],[0.10357,0.82177,0.82437],[0.10026,0.82955,0.81389],[0.09750,0.83714,0.80342],[0.09532,0.84455,0.79299],[0.09377,0.85175,0.78264],[0.09287,0.85875,0.77240],[0.09267,0.86554,0.76230],[0.09320,0.87211,0.75237],[0.09451,0.87844,0.74265],[0.09662,0.88454,0.73316],[0.09958,0.89040,0.72393],[0.10342,0.89600,0.71500],[0.10815,0.90142,0.70599],[0.11374,0.90673,0.69651],[0.12014,0.91193,0.68660],[0.12733,0.91701,0.67627],[0.13526,0.92197,0.66556],[0.14391,0.92680,0.65448],[0.15323,0.93151,0.64308],[0.16319,0.93609,0.63137],[0.17377,0.94053,0.61938],[0.18491,0.94484,0.60713],[0.19659,0.94901,0.59466],[0.20877,0.95304,0.58199],[0.22142,0.95692,0.56914],[0.23449,0.96065,0.55614],[0.24797,0.96423,0.54303],[0.26180,0.96765,0.52981],[0.27597,0.97092,0.51653],[0.29042,0.97403,0.50321],[0.30513,0.97697,0.48987],[0.32006,0.97974,0.47654],[0.33517,0.98234,0.46325],[0.35043,0.98477,0.45002],[0.36581,0.98702,0.43688],[0.38127,0.98909,0.42386],[0.39678,0.99098,0.41098],[0.41229,0.99268,0.39826],[0.42778,0.99419,0.38575],[0.44321,0.99551,0.37345],[0.45854,0.99663,0.36140],[0.47375,0.99755,0.34963],[0.48879,0.99828,0.33816],[0.50362,0.99879,0.32701],[0.51822,0.99910,0.31622],[0.53255,0.99919,0.30581],[0.54658,0.99907,0.29581],[0.56026,0.99873,0.28623],[0.57357,0.99817,0.27712],[0.58646,0.99739,0.26849],[0.59891,0.99638,0.26038],[0.61088,0.99514,0.25280],[0.62233,0.99366,0.24579],[0.63323,0.99195,0.23937],[0.64362,0.98999,0.23356],[0.65394,0.98775,0.22835],[0.66428,0.98524,0.22370],[0.67462,0.98246,0.21960],[0.68494,0.97941,0.21602],[0.69525,0.97610,0.21294],[0.70553,0.97255,0.21032],[0.71577,0.96875,0.20815],[0.72596,0.96470,0.20640],[0.73610,0.96043,0.20504],[0.74617,0.95593,0.20406],[0.75617,0.95121,0.20343],[0.76608,0.94627,0.20311],[0.77591,0.94113,0.20310],[0.78563,0.93579,0.20336],[0.79524,0.93025,0.20386],[0.80473,0.92452,0.20459],[0.81410,0.91861,0.20552],[0.82333,0.91253,0.20663],[0.83241,0.90627,0.20788],[0.84133,0.89986,0.20926],[0.85010,0.89328,0.21074],[0.85868,0.88655,0.21230],[0.86709,0.87968,0.21391],[0.87530,0.87267,0.21555],[0.88331,0.86553,0.21719],[0.89112,0.85826,0.21880],[0.89870,0.85087,0.22038],[0.90605,0.84337,0.22188],[0.91317,0.83576,0.22328],[0.92004,0.82806,0.22456],[0.92666,0.82025,0.22570],[0.93301,0.81236,0.22667],[0.93909,0.80439,0.22744],[0.94489,0.79634,0.22800],[0.95039,0.78823,0.22831],[0.95560,0.78005,0.22836],[0.96049,0.77181,0.22811],[0.96507,0.76352,0.22754],[0.96931,0.75519,0.22663],[0.97323,0.74682,0.22536],[0.97679,0.73842,0.22369],[0.98000,0.73000,0.22161],[0.98289,0.72140,0.21918],[0.98549,0.71250,0.21650],[0.98781,0.70330,0.21358],[0.98986,0.69382,0.21043],[0.99163,0.68408,0.20706],[0.99314,0.67408,0.20348],[0.99438,0.66386,0.19971],[0.99535,0.65341,0.19577],[0.99607,0.64277,0.19165],[0.99654,0.63193,0.18738],[0.99675,0.62093,0.18297],[0.99672,0.60977,0.17842],[0.99644,0.59846,0.17376],[0.99593,0.58703,0.16899],[0.99517,0.57549,0.16412],[0.99419,0.56386,0.15918],[0.99297,0.55214,0.15417],[0.99153,0.54036,0.14910],[0.98987,0.52854,0.14398],[0.98799,0.51667,0.13883],[0.98590,0.50479,0.13367],[0.98360,0.49291,0.12849],[0.98108,0.48104,0.12332],[0.97837,0.46920,0.11817],[0.97545,0.45740,0.11305],[0.97234,0.44565,0.10797],[0.96904,0.43399,0.10294],[0.96555,0.42241,0.09798],[0.96187,0.41093,0.09310],[0.95801,0.39958,0.08831],[0.95398,0.38836,0.08362],[0.94977,0.37729,0.07905],[0.94538,0.36638,0.07461],[0.94084,0.35566,0.07031],[0.93612,0.34513,0.06616],[0.93125,0.33482,0.06218],[0.92623,0.32473,0.05837],[0.92105,0.31489,0.05475],[0.91572,0.30530,0.05134],[0.91024,0.29599,0.04814],[0.90463,0.28696,0.04516],[0.89888,0.27824,0.04243],[0.89298,0.26981,0.03993],[0.88691,0.26152,0.03753],[0.88066,0.25334,0.03521],[0.87422,0.24526,0.03297],[0.86760,0.23730,0.03082],[0.86079,0.22945,0.02875],[0.85380,0.22170,0.02677],[0.84662,0.21407,0.02487],[0.83926,0.20654,0.02305],[0.83172,0.19912,0.02131],[0.82399,0.19182,0.01966],[0.81608,0.18462,0.01809],[0.80799,0.17753,0.01660],[0.79971,0.17055,0.01520],[0.79125,0.16368,0.01387],[0.78260,0.15693,0.01264],[0.77377,0.15028,0.01148],[0.76476,0.14374,0.01041],[0.75556,0.13731,0.00942],[0.74617,0.13098,0.00851],[0.73661,0.12477,0.00769],[0.72686,0.11867,0.00695],[0.71692,0.11268,0.00629],[0.70680,0.10680,0.00571],[0.69650,0.10102,0.00522],[0.68602,0.09536,0.00481],[0.67535,0.08980,0.00449],[0.66449,0.08436,0.00424],[0.65345,0.07902,0.00408],[0.64223,0.07380,0.00401],[0.63082,0.06868,0.00401],[0.61923,0.06367,0.00410],[0.60746,0.05878,0.00427],[0.59550,0.05399,0.00453],[0.58336,0.04931,0.00486],[0.57103,0.04474,0.00529],[0.55852,0.04028,0.00579],[0.54583,0.03593,0.00638],[0.53295,0.03169,0.00705],[0.51989,0.02756,0.00780],[0.50664,0.02354,0.00863],[0.49321,0.01963,0.00955],[0.47960,0.01583,0.01055]])
matplotlib.cm.register_cmap(name='turbo', cmap=matplotlib.colors.ListedColormap(turbo_colormap_data))

def calc_discernibility(cmap):
    cspaces = {
        'norm': 'sRGB1'
    }
    
    for cvd in ["deuteranomaly", "protanomaly", "tritanomaly"]:
        cspaces[cvd[:4]] = {
            "name": "sRGB1+CVD",
            "cvd_type": cvd,
            "severity": 100
        }
        
    deltas = {}
    for name, space in cspaces.items():
        # convert to sRGB1 and clip to sRGB1 representable colors
        cmap = colorspacious.cspace_convert(cmap, space, 'sRGB1')
        cmap = np.clip(cmap, 0, 1)
        
        deltas[name] = colorspacious.deltaE(cmap[:-1], cmap[1:],
                                            input_space='sRGB1',
                                            uniform_space='CAM02-UCS'
                                            )
    return deltas
```

In [4]:

```
def plot_discernibility(cmap, cyclic=False):
    """
    Modified from Matthew Petroff (License: CC0)
    https://mpetroff.net/wp-content/uploads/2019/08/colormap-discernibility.ipynb
    """
    cmap_vals = matplotlib.cm.get_cmap(cmap)(np.arange(256))[:, :3]
    deltas = calc_discernibility(cmap_vals)
    
    print("vision arclength rms")
    for k, v in deltas.items():
        print(k, np.sum(v), np.std(v * v.size))
        
    # Color cycle was created using https://colorcyclepicker.mpetroff.net/, with colors picked to match deficient cones
    colors = {'norm': '#4e4e4e', 'deut': '#99e411', 'prot': '#fb8080', 'trit': '#4eacf5'}
    # lines = {'norm': '-', 'deut': '--', 'prot': ':', 'trit': '-.'}
    
    for i, t in enumerate(deltas):
        plt.plot(deltas[t], label=t.title(), c=colors[t]) #, ls=lines[t], zorder=-i)

    plt.ylabel('deltaE')
    plt.xlim(0, 255)
    plt.ylim(0, 3)
    plt.legend(ncol=4, loc='upper center', frameon=False)
    
    ax = plt.gca().axes
    ax.get_xaxis().set_visible(False)
    plt.gca().axes.get_xaxis().set_visible(False)
    
    c = matplotlib.cm.ScalarMappable(cmap=cmap)
    c.set_array(np.linspace(0, 1, 100))
    plt.colorbar(c, orientation='horizontal', pad=0, ticks=[])
    
    plt.tight_layout()
```

##### Rainbow colormaps¶

In [5]:

```
plot_discernibility('jet')
plt.savefig('jet.pdf')
```

```
vision arclength rms
norm 237.6489067987751 102.01315608719575
deut 197.22162547245688 113.06641321510008
prot 191.30976723352893 109.79410761303582
trit 159.0088438553147 110.12141135919923
```

In [6]:

```
plot_discernibility('jet_deut')
```

```
vision arclength rms
norm 197.22162547245688 113.06641321510008
deut 191.13723626657634 123.87415018637341
prot 186.69125118671104 119.72662744985995
trit 155.16883944144877 118.03150000716025
```

In [7]:

```
plot_discernibility('turbo')
```

```
vision arclength rms
norm 229.5592316392312 82.43735042653213
deut 187.35476417648886 103.14394426706467
prot 186.76786084495728 103.6007870253746
trit 160.0383500614337 102.87039993059278
```

##### Perceptually uniform linear colormaps¶

In [8]:

```
plot_discernibility('viridis')
plt.savefig('viridis.pdf')
```

```
vision arclength rms
norm 123.87240424869012 1.4426561486446652
deut 102.06985469284778 24.75144098279799
prot 100.44040165686378 25.05074184657724
trit 83.05194394045634 15.968495175087352
```

In [9]:

```
plot_discernibility('viridis_deut')
```

```
vision arclength rms
norm 102.06985469284778 24.75144098279799
deut 102.6694597188469 26.66952824377542
prot 100.80714002872865 26.1941064958626
trit 83.94959249243949 18.102923640267903
```

In [10]:

```
plot_discernibility('cividis')
```

```
vision arclength rms
norm 99.50610470446995 5.4733668647969225
deut 100.16766444106281 10.280564146393903
prot 98.09574746453897 10.610630221875478
trit 81.95403722703333 19.40071467924089
```

##### Diverging colormaps¶

In [11]:

```
plot_discernibility('coolwarm')
```

```
vision arclength rms
norm 134.15553839484437 26.372894082222626
deut 124.40772190825527 27.104045223305032
prot 121.21768635679717 24.800709464920285
trit 106.75155412290356 24.557040448747724
```

In [12]:

```
plot_discernibility('cet_bjy')
```

```
vision arclength rms
norm 98.52805156194339 22.709018490830836
deut 92.84867888205456 27.420098344812022
prot 87.23712922807093 30.05027901980894
trit 63.34173663986917 29.783689869136584
```

##### A plot of Jet in terms of RGB primaries¶

In [13]:

```
x = np.linspace(0, 1, 1000)
cmap = matplotlib.cm.get_cmap("jet")
y = [cmap(i) for i in x]
r, g, b, a = zip(*y)

lines = {'Red': '-', 'Green': '-.', 'Blue': ':'}
plt.plot(b, label='Blue', c='blue')#, ls='-')
plt.plot(g, label='Green', c='green') #, ls='-.')
plt.plot(r, label='Red', c='red') #, ls=':')

plt.ylabel('')
plt.xlim(0, x.size)
plt.ylim(-.2, 1.2)
plt.legend(ncol=4, loc='upper center', frameon=False)

ax = plt.gca().axes
ax.get_xaxis().set_visible(False)
plt.gca().axes.get_xaxis().set_visible(False)

c = matplotlib.cm.ScalarMappable(cmap=cmap)
c.set_array(np.linspace(0, 1, 100))
plt.colorbar(c, orientation='horizontal', pad=0, ticks=[])

plt.tight_layout()
plt.savefig('jet_primaries.pdf')
```

In [ ]:

In [ ]:
